# Supplement comprising of laccase and citric acid as an alternative for antibiotics – *in vitro* triggers of melanin production

**DOI:** 10.1101/179291

**Authors:** M. Chaali, J. Lecka, G. Suresh, M. Salem, S. K. Brar, L. Hernandez-Galan, J. Sévigny, A. Avalos Ramirez

## Abstract

An indiscriminate use of antibiotics in humans and animals has led to a widespread selection of antibiotic-resistant bacterial strains. A possible solution to counter this problem could be to develop alternatives that may boost the host immunity, thus reducing in the quantity and frequency of antibiotic use. In this work, for the first time, citric acid and laccase were used as extracellular inducers of melanin production in yeast cells and human cell lines. It is proposed that the formulation of laccase and citric acid together could further promote melatonin stimulated melanocyte derived melanin production. Melanization test as a probe of immunity, described in this study, is an easy and a quicker test than the other immunity tests and is statistically significant. The results showed the synergistic effect of citric acid and laccase on melanin production by the yeast cells, with significant statistical differences compared to all other tested conditions (P: 0.0005- 0.005). Laccase and citric acid together boosted melanin production after 8 days of incubation. An increase in melanin production by two colon human cells lines (Cacao-2/15 and HT-29) was observed when both laccase and citric acid were present in cell growth medium. A formulation with citric acid and laccase may prove to be an excellent alternative to reduce the antibiotic load in human and animal subjects.

**Summary statement:** This study shows, for the first time, that production of melanin in yeast and human intestinal cells is induced by extracellular addition of laccase and citric acid.

## INTRODUCTION

Antibiotics are used for the treatment of infectious diseases (Van Boeckel, Gandra et al. 2014). However, the emergence of bacterial resistance to antibiotics has become a public concern, therefore, their indiscriminate use is imprudent and unjustified. Thus, antibiotics no longer remain valid tools to counter the onslaught of deadly bacterial human diseases (Huyghebaert, Ducatelle et al. 2011, Toghyani, Tohidi et al. 2013). Several studies have demonstrated organic acids and enzymes as biological alternatives to antibiotic (Couto and Herrera 2006, Fallah, Kiani et al. 2013). In addition to being a favourable modifier of immunity (Chowdhury, Islam et al. 2009), citric acid was also demonstrated as one of the most efficient organic acids, having a strong antimicrobial activity (Dibner and Buttin 2002). Likewise, the addition of enzymes reduced the number of gut bacteria leading to improved gut performance (Bedford 2000). Laccase is well known for its high ability to degrade the phenolic compounds (Mayer and Staples 2002) and for antioxidant activity (Prasetyo, Kudanga et al. 2009). Additionally, it was reported that laccase triggered the production of melanin (Mayer and Staples 2002, Zhu and Williamson 2004), a black pigment serving as cellular virulence against bacteria in form of melanophores (Nosanchuk, Stark et al. 2015). Melanogenesis provides protection from environmental stress conditions to various groups of free living organisms (Plonka and Grabacka 2006). In humans, melanin was shown to play an important role in intestinal inflammatory processes via melanin-concentrating hormone (MCH) (Kokkotou, Moss et al. 2008, Ziogas, Karagiannis et al. 2014). It was also reported that there is a link between melanin based pigmentary process and immune systems in the middle ear (Fritz, Roehm et al. 2014) and with inflammation accompanied obesity (Page, Chandhoke et al. 2011) in human, while in birds, melanin based coloration variations are suggested to be indicators of environmental oxidative stress in a sexually selected manner (Solano 2014). The key enzyme for melanin synthesis is tyrosinase (EC 1.14.18.1), which is also responsible for biosynthesis of dopaquinine from tyrosine in sequential reactions. Melanin is synthesized by different organisms, such as bacteria, fungi, and plants (Swan 1974). However, studies by Roy et al (Roy, Nayak et al. 1989) had reported laccase activity in yeast for the first time. In mammals, the production of melanin by melanocytes is melatonin dependent. In this study, exogenous application of laccase and citric acid on yeast has been confirmed for aggressive production of melanin in melanocytes and melatonin in an independent manner.

## MATERIAL AND METHODS

### Production of laccase

Laccase was produced as described by Gassara et al. (Gassara, Brar et al. 2010) and is briefly explained below.

### Microorganism

*Trametes versicolor* was grown in potato dextrose broth (PDB) medium in incubator shaker at 30±1°C, 150 rpm for 7 days before incubating it on potato dextrose agar (PDA) petri plate in static incubator at 30±1°C for 10 days (Lorenzo, Moldes et al. 2002).

### Solid state fermentation (SSF)

Forty grams of apple pomace (AP) was used as substrate for the SSF. The moisture was adjusted to 75% (v/w) and Tween-80 (3 mmol/kg of dry substrate) was used as an inducer for the production of laccase. After sterilization at 121 °C ±1 °C for 30 min, the substrate was inoculated with fungal mats from the PDA petri plate. The fermentation was carried out in static incubator at 30 °C ±1 °C for 15 days. The profiling of laccase production was carried out by daily sampling for analysis (Gassara, Brar et al. 2010).

### Enzyme extraction and activity test

About 50 mM Sodium-phosphate buffer (10/1, v/w) at pH 6.5 was added to one gram of sample in incubator shaker at 150 rpm for 2 h at 30 °C±1 °C.

The mixture was centrifuged at 7000 x g for 20 min, the supernatant was collected and the enzyme activity was determined spectrophotometrically at 436 nm using 2,2-azino bis(3-ethylbenzthiazoline-6-sulfonic acid) (ABTS) in 0.1 M phosphate-citrate buffer at pH 4 and 45±1°C (Gassara, Brar et al. 2010).

### Production of citric acid

The production was based on the method developed by Dhillon et al. (Dhillon, Brar et al. 2013) as explained below.

### Microorganism and spore solution

*Aspergillus niger* (NRRL 567 and NRRL 2001) was used for the production of citric acid. The fungus was grown in potato dextrose broth (PDB) medium for 6-7 days in shaking incubator followed by its incubation on PDA plate for 5 days in static incubator at 30 °C. The spores were collected from the PDA plate using a 0.1% (v/v) Tween-80 solution. Spores were counted using haemocytometer (Dhillon, Brar et al. 2011, Dhillon, Brar et al. 2013).

### Solid state fermentation (SSF)

Fifty grams of AP was used as substrate. The moisture was adjusted to 75% (v/w). The substrate was autoclaved at 121 °C±1 °C for 30 min. About 3% (v/w dry) methanol was filtered via Whatman paper and added to substrate to enhance the production of citric acid. After thorough mixing, 1×10^7^ spores/g dry substrate was added to inoculate the medium. The fermentation was performed in static incubator at 30 °C ±1 °C for 5 days. Each day, a sample was taken for the profiling studies (Dhillon, Brar et al. 2011, Dhillon, Brar et al. 2013).

### Citric acid extraction and analysis

Citric acid extraction was done by adding distilled water to substrate (solid/liquid ratio 1:15). The mixture was incubated at 30°C ±1 °C, 150 rpm for 30 min followed by centrifugation for 30 min at 7000 × *g*. The supernatant was retained for citric acid estimation measured using the Marier and Boulet method (Marier and Boulet 1958, Dhillon, Brar et al. 2011, Dhillon, Brar et al. 2013).

### Melanization test

This test was adapted to the method used by Wang et al. and Sabiiti et al., please see details below (Wang, Aisen et al. 1996, Sabiiti, Robertson et al. 2014).

#### - Yeast cells culture

For the melanization test, the yeast strain, *Saccharomyces cerevisae,* was grown on yeast broth of peptone dextrose (YPD) agar cultured in 3 ml YPD broth in shaker incubator at 20 rpm, for 24 h at 25 °C ±1 °C. The yeast cells were counted using a haemocytometer (Wang, Aisen et al. 1996, Sabiiti, Robertson et al. 2014).

#### - Test conditions

The medium was prepared by adding to 75 mL distilled water: 15 mM glucose, 10 mM MgSO4, 29.4 mM KH2PO4, 13 mM glycine. The medium was sterilized at 121 °C ±1 °C for 30 min, after which 3.0 mM vitamin B1 and 1.0 mM catechol was added under aseptic conditions (Wang, Aisen et al. 1996, Sabiiti, Robertson et al. 2014).

The test was performed thrice under four conditions with the number of the cells being the same: A: control with only yeast cells (1.0 × 10^8^ yeast cells), B: yeast cells and citric acid (15g/L), C: yeast cells and laccase (1400 U) and; D: yeast cells with laccase (1400 and citric acid (15g/L). Each condition was represented in triplicates. The flasks were incubated 150 rpm at 30 °C ±1°C, for 10 days (Wang, Aisen et al. 1996, Sabiiti, Robertson et al. 2014). A kinetic study of melanin production was done by taking a sample every two day.

#### - Yeast cell count

Yeast cells were centrifuged at 9000 *× g* for 20 min. The pellet was washed with 1.0 M sorbitol in 0.1 M sodium citrate (pH 5.0), and resuspended in 5 mL of the same solution. The cells were counted using haemocytometer (Wang, Aisen et al. 1996, Sabiiti, Robertson et al. 2014).

#### - Human colon carcinoma cells culture and melanization test

HT-29 (ATCC® HTB-38TM) human colon adenocarcinoma cell line was purchased from the American Type Culture Collection and the human colon carcinoma cell lines Caco-2/15 were provided by J. F. Beaulieu, Université de Sherbrooke, Sherbrooke, Québec, Canada. Both cell lines were maintained in monolayer cultures in DMEM/F-12 growth medium supplemented with Glutamax (2 mM), antibiotic-antimycotic solution (1X), Hepes (25 mM), Normocin (used as an antimycoplasma reagent, 100 μg/mL), and 10% heat-inactivated fetal bovine serum at 37°C in a 95% air: 5% CO2 atmosphere. Cells were regularly monitored for the presence of *Mycoplasma* spp. by means of a conventional PCR test (Wirth, Berthold et al. 1994) using 5 μg of extracted genomic DNA (PureLink genomic DNA mini kit) as a template. The cells from passages 2-3 were seeded (1.5 × 106/well) in 6-well plates containing 3 mL medium and 3 mL of melanization solution (30 mM glucose, 20 mM MgSO4, 58.8 mM KH2PO4, 26 mM glycine, 6.0 mM vitamin B1, 2.0 mM catechol, 2800 U laccase and 30g/L citric acid). After 3 days, cells and media were collected for melanin quantification (similar to yeast cells).

#### - Melanin extraction and quantification

The cells were sonicated for 45 seconds. The cell debris was collected by centrifugation at 9000 × g for 20 min and resuspended in 4.0 M urea for 1h at room temperature. The cell debris was collected by centrifugation at 9000 x *g* for 20 min, and resuspended in 6.0 M HCl at 100°C for 1h. The solution was then filtered through Whatman paper (45μm) using a vacuum pump. The filter was dried completely in a desiccator before weighing to quantify melanin produced by the yeast cells by adapting the method used by Wang et al. and Sabiiti et al. (Wang, Aisen et al. 1996, Sabiiti, Robertson et al. 2014).

### Comparison using FTIR

The melanin produced by the cells was taken from filters and dissolved in 2M NaOH solution. The same was done for the pure melanin standard, purchased from Sigma-Aldrich. Both solutions were analyzed using FTIR (Fourier Transform Infrared Spectroscopy). The wavelengths and intensities of the two spectra obtained by the FTIR are presented using Excel.

### Statistical analysis

Statistical analysis was done using Variance (ANOVA) by Bonferroni’s multiple comparisons test (Prism software) and Student’s *t*-test. *P*-values below 0.05 were considered statistically significant.

## RESULTS

*Trametes versicolor* was selected for its high potential of production of laccase. The highest enzyme production at 35 U/g _dry substrate_ was obtained in 13 days of incubation at 30° C. (Fig.1). *Aspergillus niger* was chosen for its copious citric acid production capacity, Fig. 2 presents profile of citric acid production. Two peaks were obtained between 40 and 80 h of incubation. The highest citric acid production averaged at 35.4±0.7 g/kg dry substrate.

**Fig. 1:**
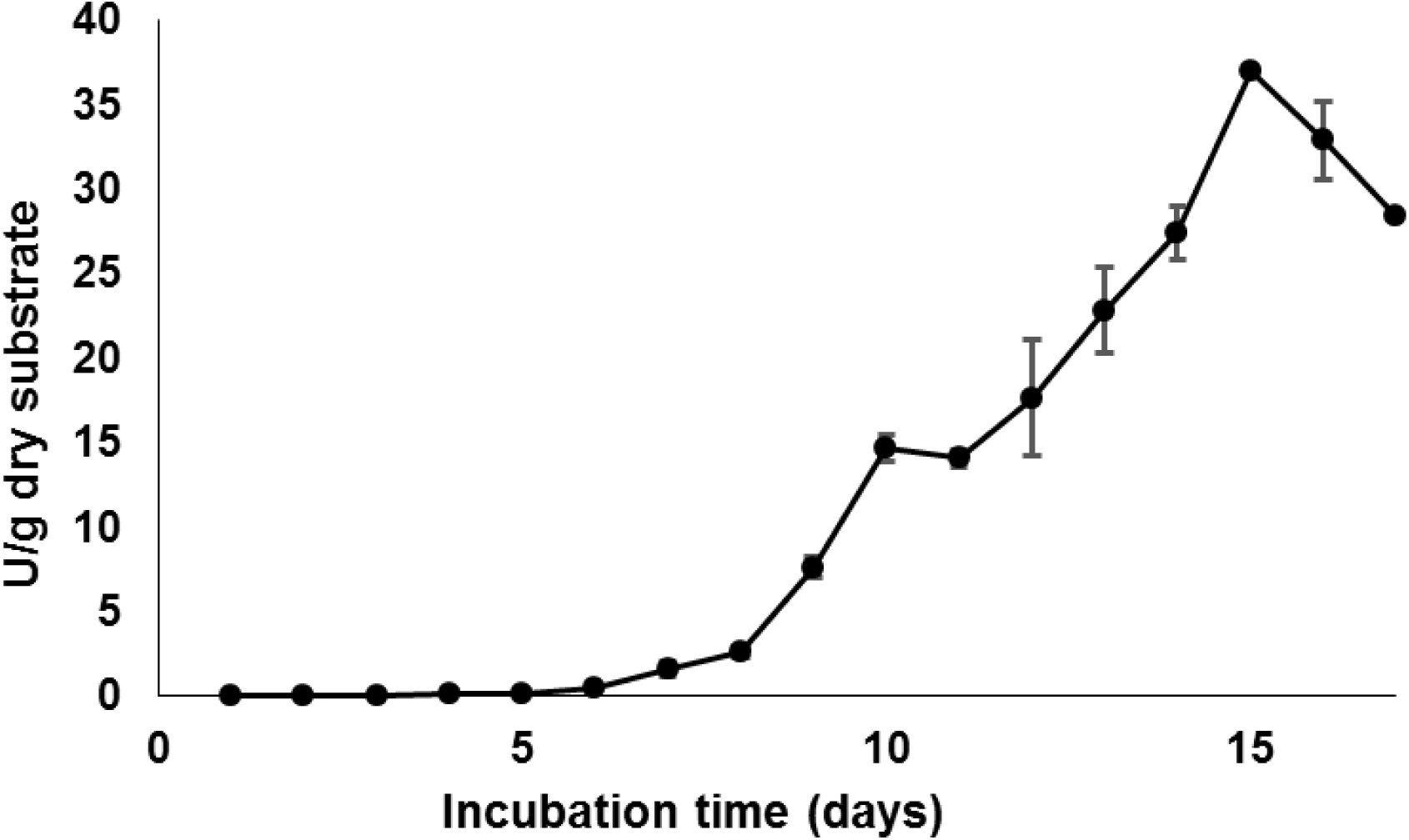
Laccase production by *T. versicolor* in solid state fermentation using apple pomace as substrate. The graph represents mean values ±SD (n=6).

**Fig. 2:**
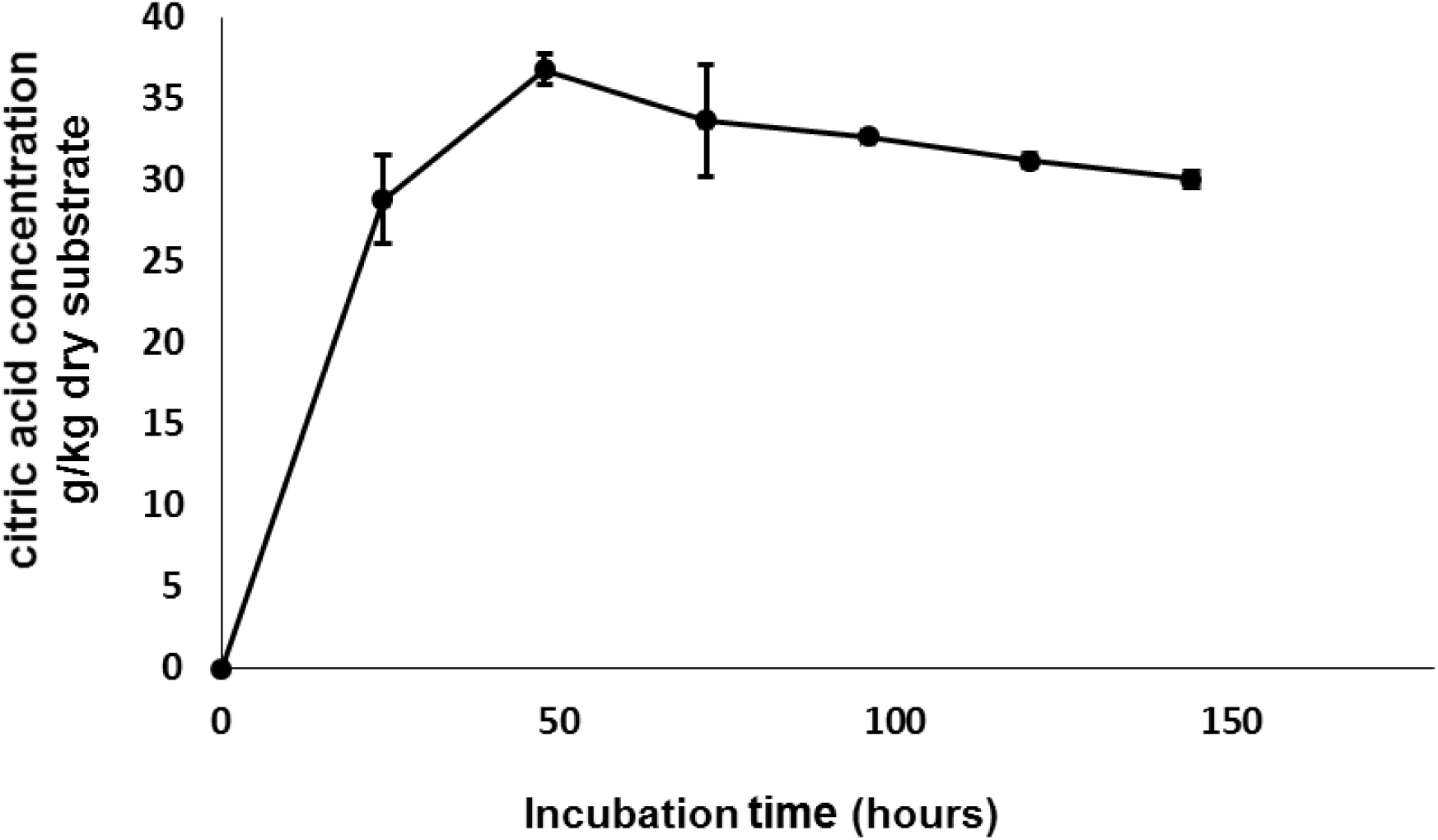
Citric acid production by *A. niger* in solid state fermentation using apple pomace as substrate. The graph represent mean values ±SD (n=6).

For the melanization tests, the yeast cells were counted for each condition after 10 days of incubation. Melanin from yeast cells was extracted using sonication and weighed. The cell count was similar for all the conditions, however, significant differences were observed in melanin harvested from yeast cells raised with laccase (Table1). The highest production of melanin by the yeast cells was 1.04 × 10^−9^g, and was observed when both laccase and citric acid were present in the growth medium (Fig.3A, B and C).

**Table 1.** Statistical analysis of cell survival (n=10), melanin production per total yeast cells (n=10) and melanin production per cell (n=10) in different conditions during melanin test: (A) cells only (control), (B) cells in the presence of citric acid, (C) cells in the presence of laccase, (D) cells in the presence of citric acid and laccase

| Analysis of Variance (ANOVA) by Bonferroni's multiple comparisons test using Prism software. |  |  |  |  |  |  |
| --- | --- | --- | --- | --- | --- | --- |
| **P values** |  |  |  |  |  |  |
| Tested parameters | **A:B** | **A:C** | **A:D** | **B:C** | **B:D** | **C:D** |
| cell survival | **0.9999 (ns)\*** | **0.009** | **0.0078** | **0.2904 (ns)\*** | **0.2406 (ns)\*** | **0.9999 (ns)\*** |
| melanin production per total cell number | **0.99 (ns)\*** | **0.1965 (ns)\*** | **0.1991 (ns)\*** | **0.4131 (ns)\*** | **0.4108 (ns)\*** | **0.9999 (ns)\*** |
| melanin production per cell | **0.9999 (ns)\*** | **0.0002** | **0.0001** | **0.0016** | **0.0001** | **0.4749 (ns)\*** |

\*ns: statistically non-significant

**Fig. 3A:**
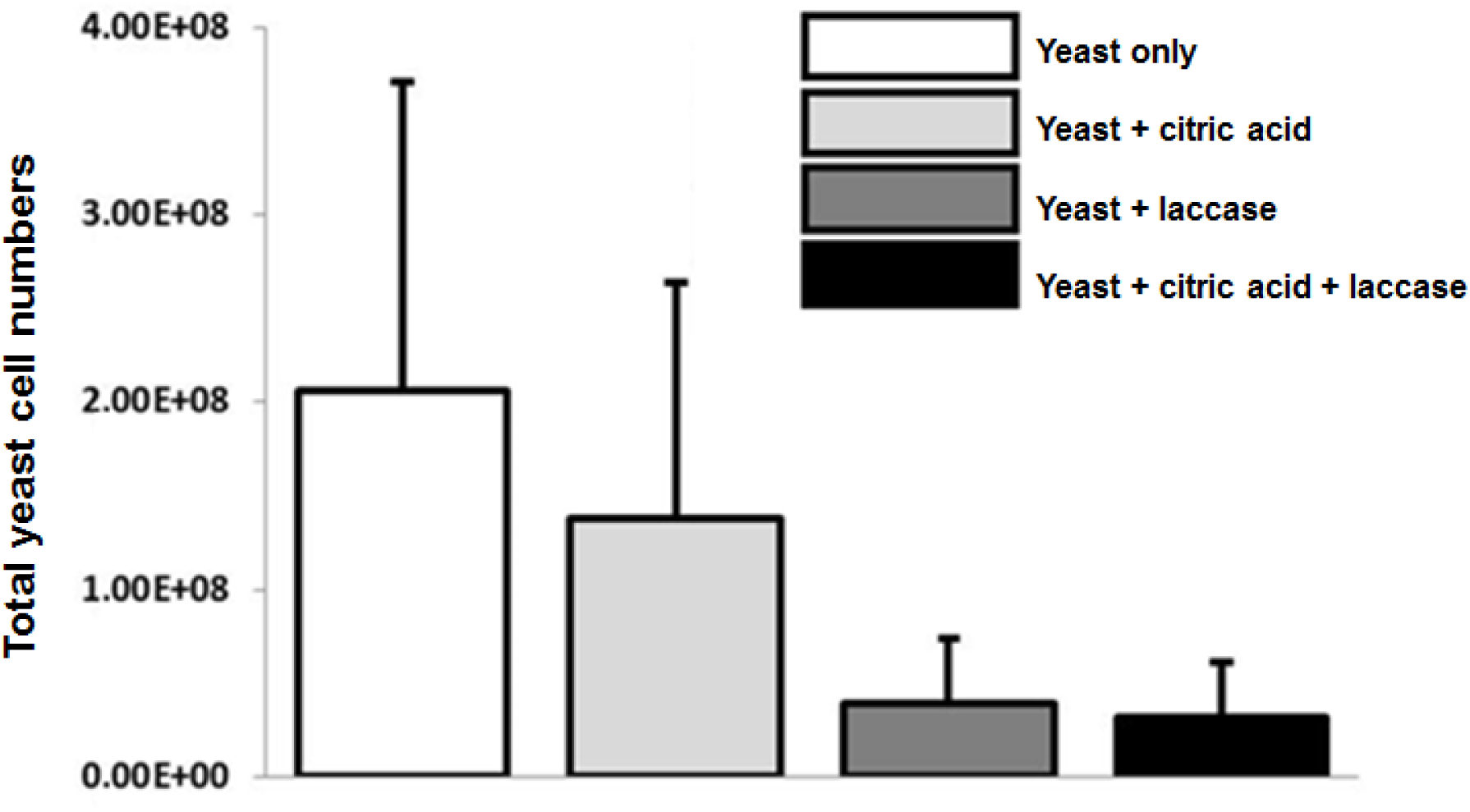
Total yeast cell count after 10 days of incubation under different test conditions. The graph shows the mean values ±SD (n=20).

**Fig. 3B:**
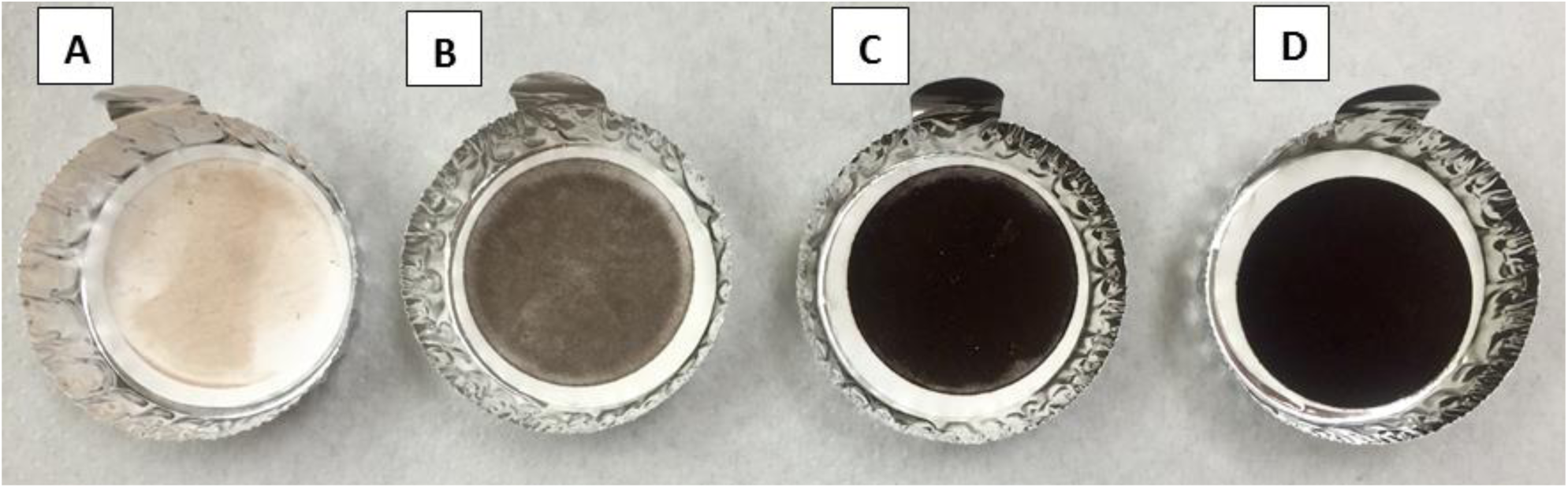
Melanin production by: (A) cells only (control), (B) cells in presence of citric acid, (C) cells in presence of laccase, (D) cells in presence of citric acid and laccase. The photograph represent one out of 20 experiments.

**Fig. 3C:**
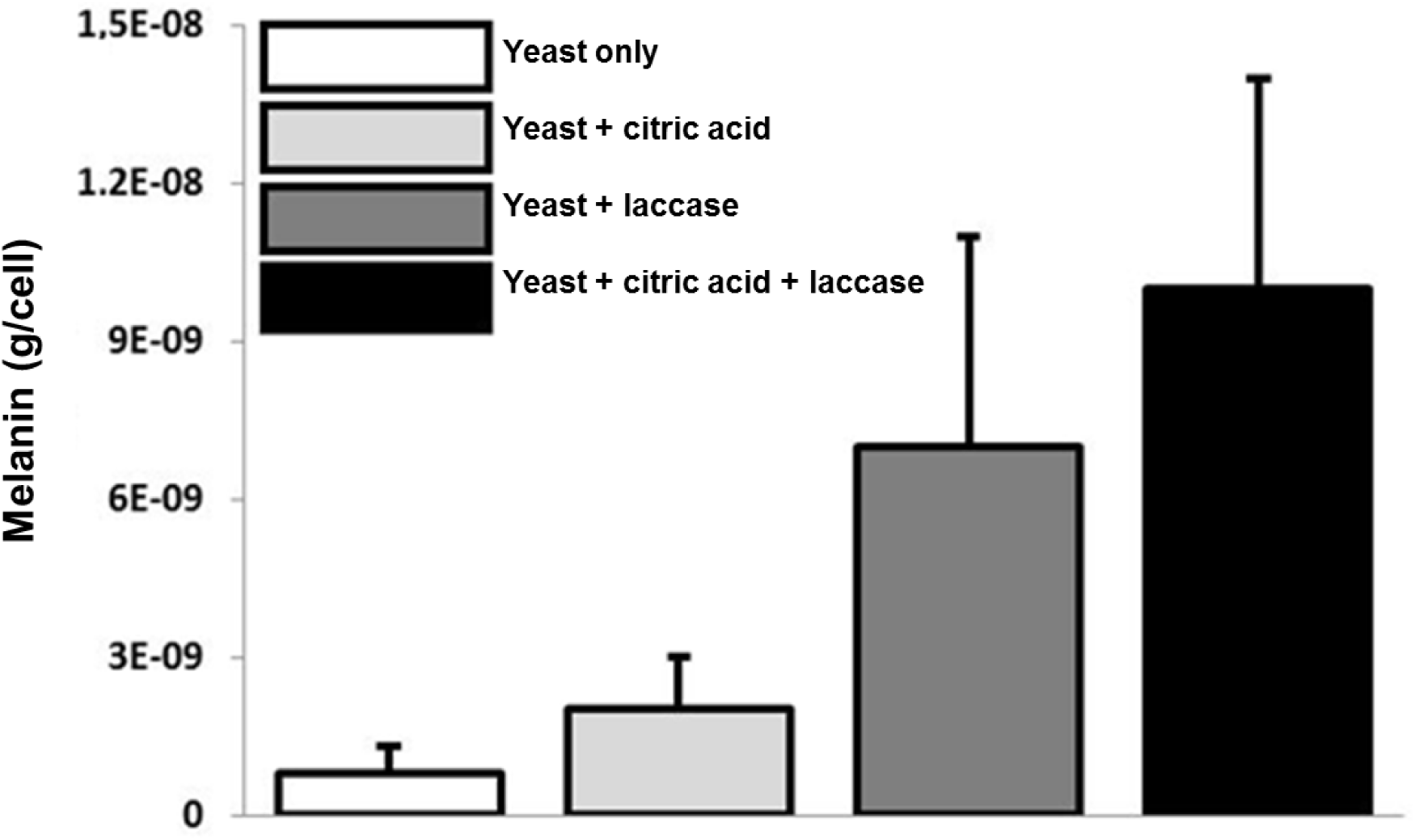
Melanin production per yeast cell after 10 days of incubation under different test conditions. The graph shows the mean values ±SD (n=20).

Kinetic studies of melanin production by yeast cells is presented in Fig.4, for which sampling was done every two days up to day 10^th^. During first 6 days, melanin production was slow and the production increased significantly (P: 0.02-0.003) after 8 days of incubation (3.399 x10^-9^ g/cell) (Table.2).

**Fig. 4:**
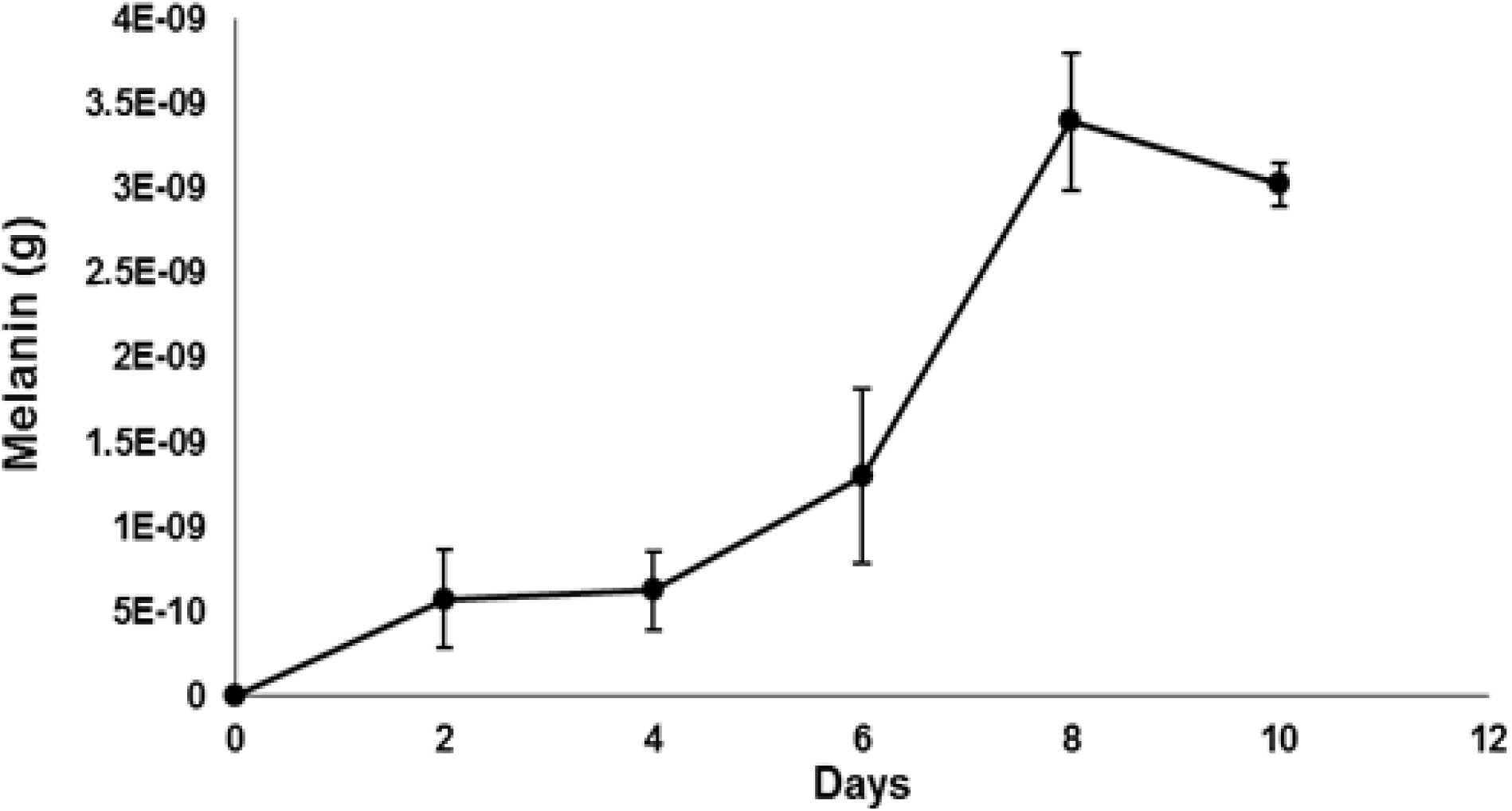
Melanin production kinetics by yeast cells in the presence of citric acid and laccase. The graph represents one out of 20 experiments

**Table 2.**
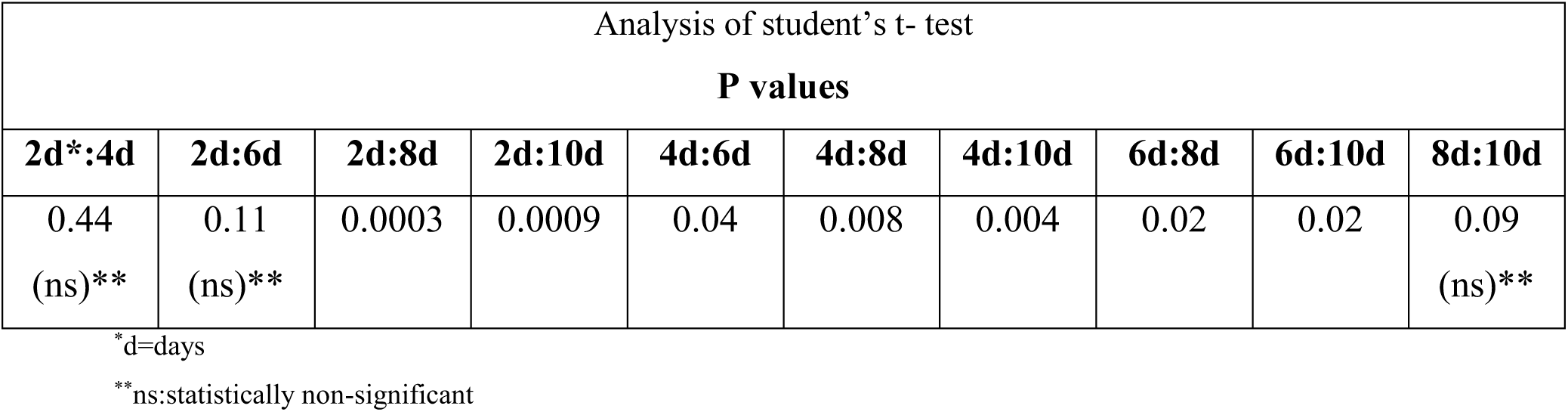
Statistical analysis of melanin production per yeast cells (n=3) in the presence of citric acid and laccase in different time period (2-10 days)

| Analysis of student's t- test |  |  |  |  |  |  |  |  |  |
| --- | --- | --- | --- | --- | --- | --- | --- | --- | --- |
| **P values** |  |  |  |  |  |  |  |  |  |
| **2d\*:4d** | **2d:6d** | **2d:8d** | **2d:10d** | **4d:6d** | **4d:8d** | **4d:10d** | **6d:8d** | **6d:10d** | **8d:10d** |
| 0.44<br>(ns)** | 0.11<br>(ns)** | 0.0003 | 0.0009 | 0.04 | 0.008 | 0.004 | 0.02 | 0.02 | 0.09<br>(ns)** |

\*d=days
\*\*ns:statistically non-significant

Further, to observe extracellular laccase and citric acid induced melanin production in human cells, they were studied with two different cell lines of human colon carcinoma: Caco-2/15 and HT-29 and qualitative production of melanin was noted as given in Fig. 5. In human cell lines, laccase and citric acid induced increase in melanin production was rather higher than melanin produced by the yeast cells under similar condition just in 3 days (6.40x10^-9^g/cell Caco-2/15, 1.08x10^-8^g/cell HT-29 and 6.25x10^-10^g/yeast cell).

**Fig. 5:**
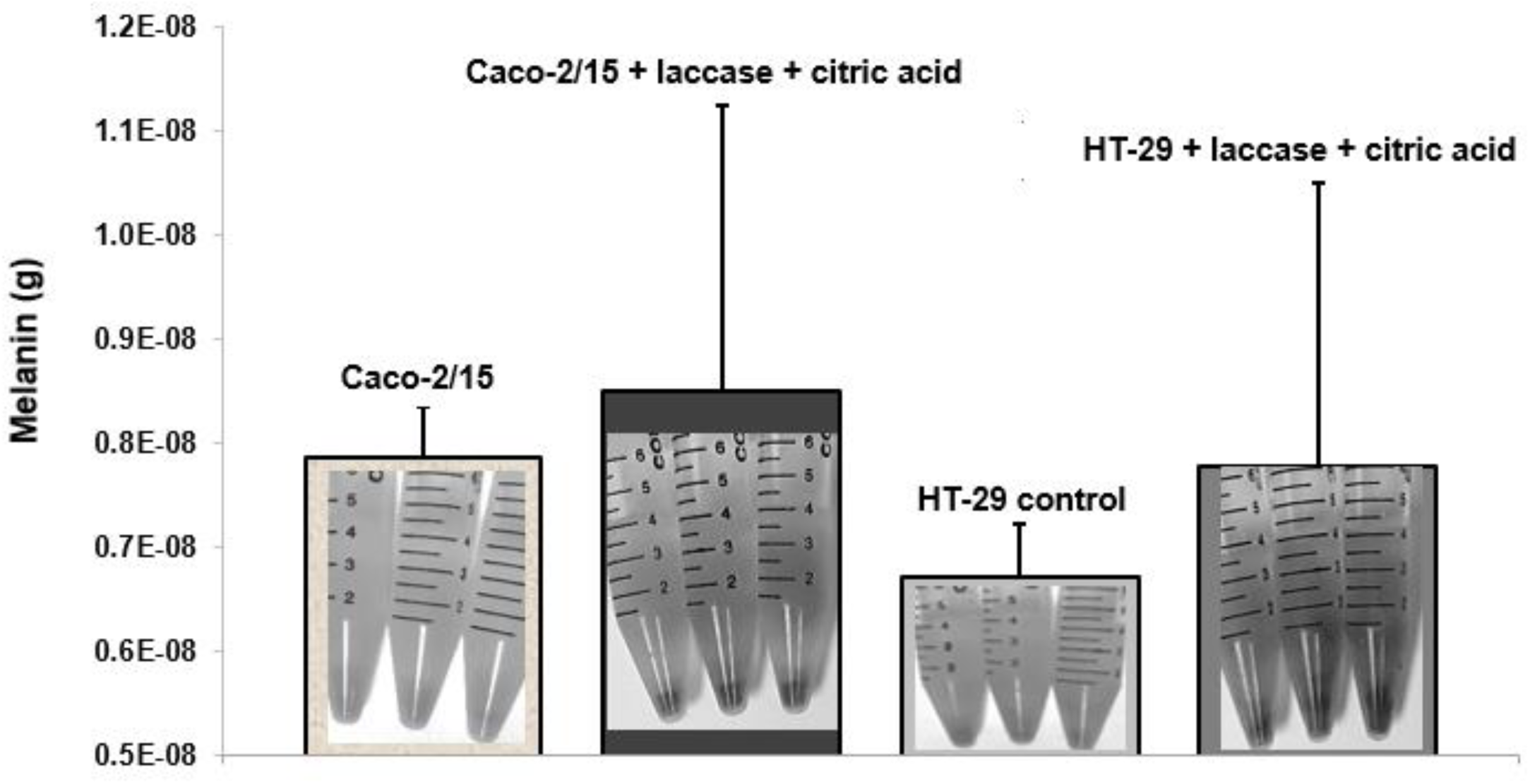
Melanin production by human colon carcinoma cells (Caco-2/15 and HT-29) after 3 days of incubation by addition of laccase and citric acid. The graph shows the mean values ±SD (n=3).

The FT-IR (compound bounds analysis) for yeast produced melanin and standard melanin showed almost the same spectra (Fig. 6) confirming identity of the produced compound.

**Fig.6:**
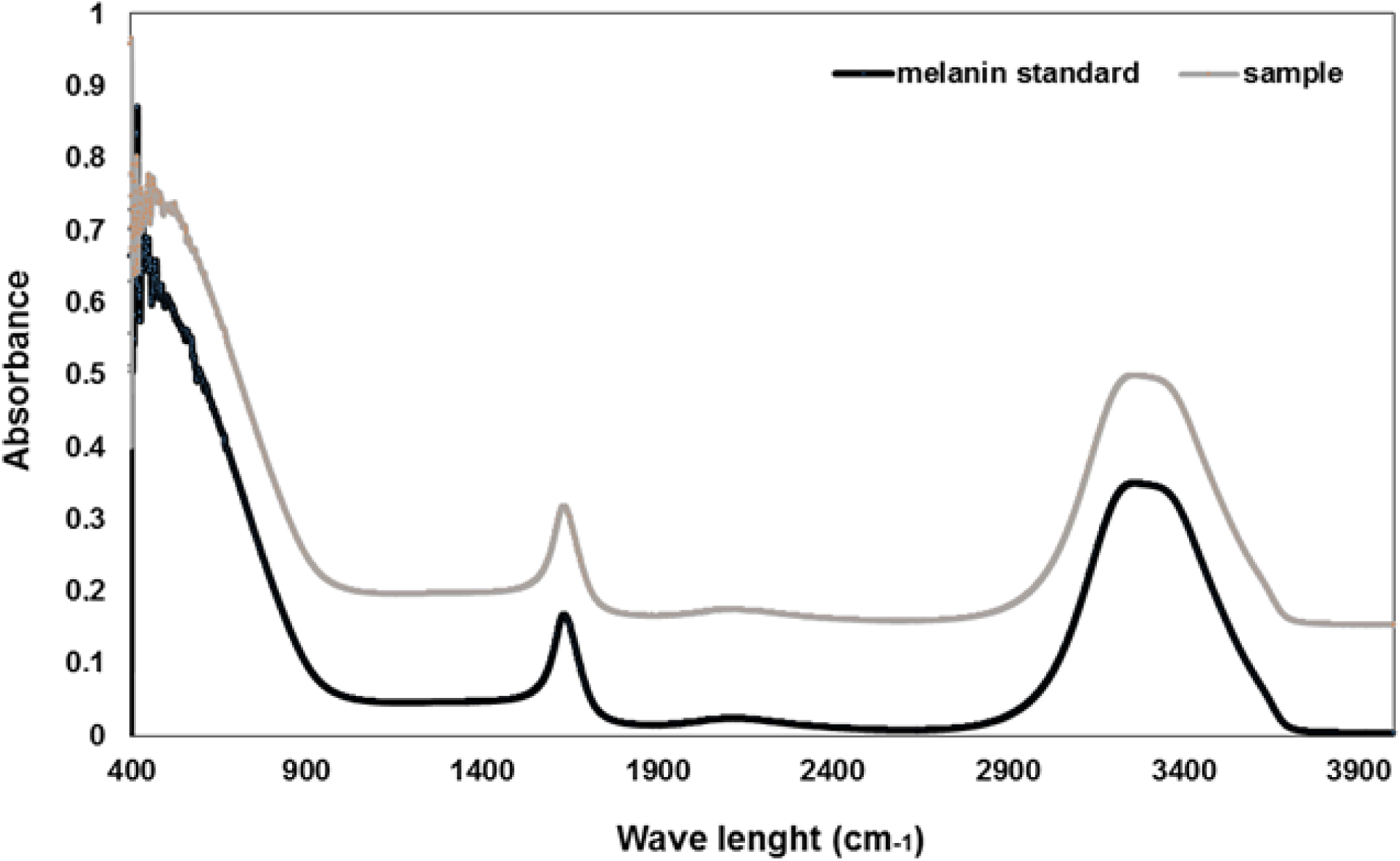
FTIR spectrum comparison between melanin standard and melanin produced by yeast cells.

## DISCUSSION

Melanin production by yeast system was used to test the efficacy of a formulation comprising fungal laccase and citric acid.

The results suggested that this formulation as a feed supplement was an eco-friendly green alternative of antibiotics. (Solano 2014) reported that phenolases (copper-proteins like laccases can oxidize phenols for this study) are characteristic enzymes related to melanin production. It is also implicated in synthesis of DHN-melanin and pyomelanin, the microbial melanins. In fungal cells, laccase acts as an anti-oxidant via the production of eumelanin, a “good” form of melanin (Williamson, Tomlinson et al. 1997, Zhu, Gibbons et al. 2001). The fact that mammals do not produce laccase shows this enzyme as of potential interest for pharmacological studies. The data here showed that after incubation, the yeast cell count differed significantly from the control group only when laccase was present in the growth media (Fig. 3A and Table 1). Similar melanomas cell number also depended on citric acid doses (Bhatnagar, Srirangam et al. 1998). Nevertheless, citric acid as a diet supplement in young broiler chickens breeding regimens was shown to be promoting their growth by increasing gastric proteolysis (Salgado-Tránsito, Del Río-García et al. 2011). It was also shown that organic acids were also important contributors to human large intestinal function (Sabboh-Jourdan, Valla et al. 2011).

The data demonstrated that laccase from innocuous yeast could mimic the melatonin action of production of melanin, an antibacterial trigger. It was already shown that mice infected with fungus with high laccase activity exhibited strong inflammatory protection (Steenbergen, Shuman et al. 2001).

Hence, melanin (black colored) extracted from each sample filtered on Whatmann paper by vacuum filtration was examined (Fig 4A): panel A represents control, B the presence of citric acid, C presence of laccase and, D that of laccase and citric acid together. Compared with other conditions, citric acid and laccase together in yeast growth medium showed synergistic effect for 10 days. The quantity of melanin produced by the cells was significantly higher, with P values between 0.0005 and 0.005 (Fig. 4B; Table 1).

The existing reports demonstrate that intracellular fungal laccase possess the ability to biosynthesize the melanin and that mutant yeast cells with higher laccase activity produced increased quantity of melanin (Zhu, Gibbons et al. 2001, Sabiiti, Robertson et al. 2014). To the best of our knowledge, this is a first study of its kind that showed that adding laccase exogenously induced yeast cells and human intestinal cells (Fig. 5, Table 4) to produce melanin and that together with citric acid, the cells showed strong synergistic effect on melanin production in yeast (Fig. 3B; Table 2,3). Thus, using both laccase and citric acid together as supplements could mimic and or supplement melatonin driven melanin production abating dual requirement of melatonin and antibiotics.

## CONCLUSION

In this study, the extracellular presence of laccase and citric acid in the yeast and human colon cells growth media induced intracellular production of melanin (statically significant after 8 days in yeast cells, P-values: 0.02-0.003; Table2). The observed phenomenon occurred in a melatonin independent manner. As a supplement, through their antioxidant and antibacterial properties, laccase and citric acid may mitigate the problem of bacterial antibiotic resistance that is coupled with adverse health and environmental impacts. This could have possible use in humans as well as in animal feed and could lead to a possible reduction in the use of antimicrobials.

## ACKNOWLEDGEMENTS

The authors are sincerely thankful to the Natural Sciences and Engineering Research Council of Canada (Discovery Grant), and FRQNT EQUIPE grant for financial support. The views or opinions expressed in this article are those of the authors.

### COMPETING INTERESTS

No competing interests declared.

